# Predicting and interpreting large scale mutagenesis data using analyses of protein stability and conservation

**DOI:** 10.1101/2021.06.26.450037

**Authors:** Magnus H. Høie, Matteo Cagiada, Anders Haagen Beck Frederiksen, Amelie Stein, Kresten Lindorff-Larsen

## Abstract

Understanding and predicting the functional consequences of single amino acid is central in many areas of protein science. Here we collected and analysed experimental measurements of effects of >150,000 variants in 29 proteins. We used biophysical calculations to predict changes in stability for each variant, and assessed them in light of sequence conservation. We find that the sequence analyses give more accurate prediction of variant effects than predictions of stability, and that about half of the variants that show loss of function do so due to stability effects. We construct a machine learning model to predict variant effects from protein structure and sequence alignments, and show how the two sources of information are able to support one another. Together our results show how one can leverage large-scale experimental assessments of variant effects to gain deeper and general insights into the mechanisms that cause loss of function.

## Introduction

The ability to predict and understand the effects of amino acid changes to protein structure, stability and function plays central roles in a number of areas of protein science. For example, improving protein function or stability is key in many biotechnological applications, and the ability to understand and predict loss of function is of central importance in disease biology. Large-scale human genome sequencing efforts are revealing millions of missense variants that change the amino acid sequences of proteins, but we do not yet know the functional consequences for most of these variants.

Studies of the effects of single amino-acid changes also present opportunities to test our understanding of the protein structure-function relationship, and the interplay with the cellular environment. Generally speaking, while a large fraction of single amino-acid substitutions in a given protein are relatively well tolerated, there is a subset that has significant detrimental consequences (***Schaafsma and Vihinen, 2017; Gray et al., 2017***). Pinpointing which variants are in the detrimental group, and the biochemical and biophysical mechanisms underlying loss of fitness, is important for example for assessing pathogenicity of so-called variants of *uncertain significance* (***Richards et al., 2015***) and understanding the mechanistic origins of disease.

Recent technological advances have enabled high-throughput assays that can quantify changes in activity, stability or other protein properties of interest for thousands of variants in a single experiment. The assays also provide large sets of data with systematic fitness profiles of variants, often providing both mechanistic insight and systematic assessment of computational models for predicting variant effects. Briefly, such Multiplexed Assays of Variant Effects (MAVEs, also called Deep Mutational Scanning experiments) have three components: (i) generation of a DNA library encoding a comprehensive set of protein variants to be tested, (ii) an assay that selects or screens for function or other properties of interest, and (iii) sequencing before and after selection to determine how each variant fares in the assay (***Fowler and Fields, 2014***). These experiments are enabled by advances in oligonucleotide synthesis and lowered cost of next-generation sequencing, and for example make it possible to create and probe saturation mutagenesis libraries (i.e. exhaustive generation of all 19 possible substitutions across the whole protein or region of interest). While library generation and sequencing is often transferable across systems, development of assays, however, usually needs to be individually tailored for the protein of interest and the function to be tested. Typically, the function of the protein is coupled to a phenotype that is amenable to high-throughput evaluation, such as growth or production of a fluorescent marker (***Fowler and Fields, 2014***). This enables quantification of the level of function ranging from wild-type-like, to moderately affected or total loss of function. The need for customised assays, however, also means that functional scores (sometimes called fitness scores) extracted from different MAVEs are often not directly comparable to one another without further normalization.

The data generated by MAVEs have been shown to help predict the status of pathogenic and benign variants, while also serving as useful benchmarks for computational variant classification methods (***Livesey and Marsh, 2020; Frazer et al., 2020***). More generally, the data can also provide detailed insight into general aspects of protein structure and function (***Gray et al., 2017; Ahler et al., 2019b; Dunham and Beltrao, 2020; Chiasson et al., 2020; Starr et al., 2020; Cagiada et al., 2021; Amorosi et al., 2021***). The extensive coverage of variants measured in the same assay provides a rich source of data that may be used for protein design, structure prediction and identification of crucial regions related to function and stability (***Stein et al., 2019***). For example, recent analyses of more than 30 data sets generated by MAVE point to individual amino acids performing different functions depending on their chemical environment (***Dunham and Beltrao, 2020***).

A given amino acid change may affect multiple properties or functions of a protein, and by combining different assays it may be possible to disentangle which substitutions affect which of those properties and functions. As most proteins need to be folded to function, assays probing protein stability and cellular abundance have received special attention. Thus, a specific type of MAVE termed VAMP-seq has been developed to probe cellular protein abundance (***Matreyek et al., 2018***), and been shown to correlate with measurements of protein stability (***Matreyek et al., 2018; Suiter et al., 2020***). While the detailed relationship between protein stability and abundance is complicated and not fully understood (***Hingorani and Gierasch, 2014; Stein et al., 2019***), we and others have shown that unstable proteins are often targeted for proteasomal degradation leading to lowered cellular abundance (***Nielsen et al., 2017; Chen et al., 2017; Nielsen et al., 2017; Scheller et al., 2019; Abildgaard et al., 2019; Nielsen et al., 2020***). Thus, by combining the results from a VAMP-seq experiment with a MAVE probing protein activity it is possible to distinguish between variants that cause loss of function due to lowered abundance from those that change the intrinsic activity of a protein (***Jepsen et al., 2020; Chiasson et al., 2020; Cagiada et al., 2021; Amorosi et al., 2021***). These experiments and analyses suggest that a relatively large fraction of variants that cause of loss-of-function are due to loss of stability and resulting degradation in the cell. Thus, it is not surprising that a large number of disease-causing variants are proteasomal targets and are found at low cellular levels (***Meacham et al., 2001; Yaguchi et al., 2004; Olzmann et al., 2004; Ron and Horowitz, 2005; Yang et al., 2011, 2013; Arlow et al., 2013; Nielsen et al., 2017; Chen et al., 2017; Nielsen et al., 2017; Scheller et al., 2019; Abildgaard et al., 2019; Nielsen et al., 2020***).

Although it is possible to perform multiplexed assays on multiple genes in a single experiment (***Després et al., 2020; Jun et al., 2020; Hanna et al., 2021; Cuella-Martin et al., 2021***), we are still far from able to probe all possible variants in all proteins by experiments. Thus, computational methods are important to predict and understand variant effects, and in some cases they may be even be more accurate than MAVEs for this purpose (***Jepsen et al., 2020; Frazer et al., 2020***). Computational variant prediction methods make it possible to estimate effects of variants that have never been seen and examine proteins that have not been studied in experiments. This aspect is important for applications in clinical genetics where many variants are extremely rare and may arise *de novo*. Of more practical importance, they can also bring variant effects onto a common scale comparable across the proteome.

Most variant effect predictors are based on features extracted from evolutionary conservation of homologous proteins, biophysical calculations based on structure, and general knowledge of amino-acid properties (***Yue et al., 2005; Kumar et al., 2009; Adzhubei et al., 2010; Casadio et al., 2011; De Baets et al., 2012; Kircher et al., 2014; Choi and Chan, 2015; Ioannidis et al., 2016; Ancien et al., 2018; Wagih et al., 2018; Gerasimavicius et al., 2020; Livesey and Marsh, 2020***). As some of these methods have been trained and tested on classification of clinical variants, it has been argued that comparison against data from MAVEs provides a useful and unbiased alternative to benchmark such methods (***Livesey and Marsh, 2020***). In such tests, it has been shown that various sequence-based approaches, including deep-learning methods, can achieve very high accuracy (***Riesselman et al., 2018; Livesey and Marsh, 2020***). Indeed, we and others have successfully applied sequence analysis and biophysical stability calculations for identification and analysis of pathogenic variants, although these methods were not trained on clinical variants (***Pey et al., 2007; Yin et al., 2017; Nielsen et al., 2017; Gray et al., 2018; Scheller et al., 2019; Cline et al., 2019; Abildgaard et al., 2019; Jepsen et al., 2020; Frazer et al., 2020***).

Experiments and computational analyses such as those discussed above are now beginning to provide a consistent picture of the effects of variants on protein stability and function. Variants that cause substantial loss of stability are generally found at low protein levels in the cell, and thus often lead to a loss-of-function phenotype and disease. Hence, when a variant is predicted to be highly destabilizing it is likely to be non-functional. The reverse, however, does not hold true. Variants that do not perturb protein stability may still cause loss of function via other mechanisms, such as perturbing active site residues in an enzyme or key interaction sites for binding. Such effects can often be captured by evolutionary sequence analyses which are potentially sensitive to all conserved molecular mechanisms that lead to loss of function.

Here, we aim to provide further insight into the relationship between variant effects on protein stability and function, and how computational predictions of changes in thermodynamic stability and analysis of sequence conservation may be used to predict variant effects. We use the Rosetta software to predict changes in thermodynamic stability (ΔΔG) (***Park et al., 2016; Leman et al., 2020; Frenz et al., 2020***) using as input the structure of each protein. Similarly, we use GEMME (Global Epistatic Model for predicting Mutational Effects) (***Laine et al., 2019***) to analyse multiple sequence alignments and calculate a score, which we term ΔΔE, and which captures conservation of amino acids through reconstructing phylogenetic trees. To assess their predictive power on functional variant consequences, we have collected 39 data sets previously generated by MAVEs on 29 proteins, and analyse these using the ΔΔG and ΔΔE calculations. As evolution disfavours unstable proteins (***Bloom et al., 2006; Echave and Wilke, 2017***), there is some correlation between the two approaches, however, there are also differences, e.g. for active sites, where variants are typically only identified as detrimental by evolutionary sequence analysis (***Cheng et al., 2005; Echave, 2019; Jepsen et al., 2020; Cagiada et al., 2021***). We train a machine learning model that uses ΔΔG and ΔΔE as input to predict variant effects as probed by MAVE experiments. We show that both components of the model contribute to prediction accuracy, and use the final model to provide a global view of the relationship between stability and function. In this way, our analysis of MAVE experiments using ΔΔG and ΔΔE calculations help pinpoint which variants lose function due to loss of stability. Together, our results show how loss of stability is an important contributor to loss of function, and point to future improvements for predictions of variant effects.

## Results

### Analysing Variant Effects from MAVEs by Calculations of Stability and Conservation

We first aimed to quantify how well analyses of protein stability and sequence conservation are able to capture experimental measurements of variant effects. We thus collected 39 data sets generated by MAVEs on 29 proteins from the literature (Tables S1 and S2). As we aimed to use the data in a globally-trained machine learning model we used rank normalisation to bring the original variant scores reported by the individual studies onto a common scale (*s*_exp_), where *s*_exp_ ~ 1 corresponds to wild-type-like activity in the experiment and *s*_exp_ ~ 0 corresponds to variants with low activity in the assay used in the MAVE.

For each of the 29 proteins we used Rosetta (***Park et al., 2016; Leman et al., 2020; Frenz et al., 2020***) to predict changes in thermodynamic stability (ΔΔG) for each of the 19 possible variants at each of the positions resolved in the experimental structures. Here, ΔΔG = 0 corresponds to the same stability as the wild-type protein and variants with ΔΔG > 0 correspond to those that are predicted to destabilise the protein. We also built a multiple sequence alignment for each of the proteins and analysed these using GEMME (***Laine et al., 2019***). Following rank-normalisation for each protein, the resulting GEMME ΔΔE scores quantify how likely each of the 19 substitutions are in terms of what has been observed in the evolutionary record, with ΔΔE ≈ 0 corresponding to very conservative substitutions and ΔΔE ≈ 1 corresponding to variants that are extremely rare or absent from the alignment and hence predicted to be disruptive. In total, we thus collected triplets of *s*_exp_, ΔΔG and ΔΔE for 154,808 single amino-acid variants covering 10,012 positions in the 29 proteins (Fig. 1a).

**Figure 1.**
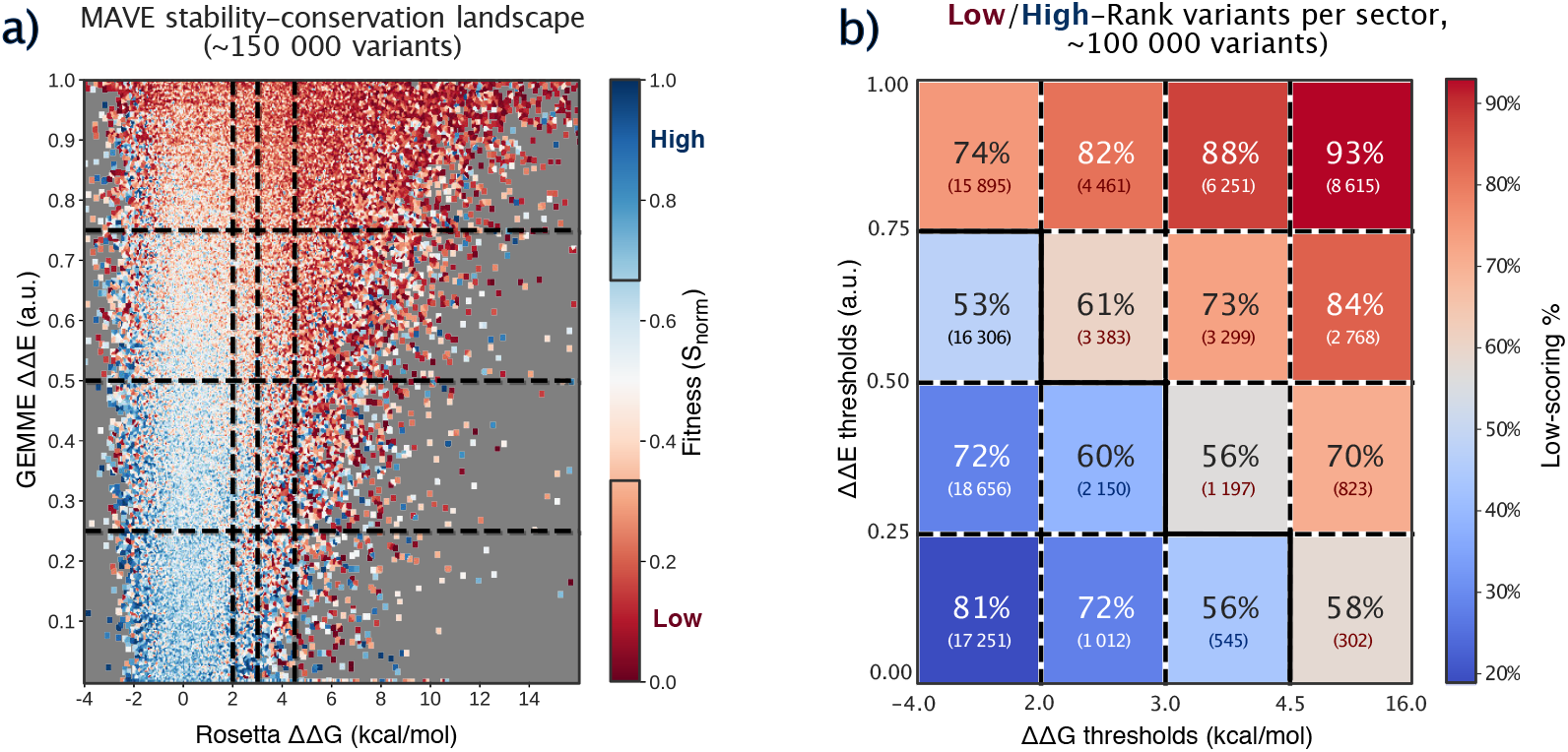
Stability and conservation score trends across variants. a) Analysis of the variant fitness landscape across over 150,000 variants in 39 MAVE experiments, by GEMME ΔΔE and Rosetta ΔΔG score per variant, colored by normalized MAVE fitness score. b) Percentage of high fitness (top 33 % of individual MAVE experimental scores, blue) or low fitness (bottom 33 %, red) variants per sector in the fitness landscape, and number of variants per sector. The middle tertile (33rd-66th percentile) of variants are excluded here.

We have previously shown that changes in protein stability(ΔΔG)and multiple-sequence-alignment-based conservation scores (ΔΔE) correlate with changes in cellular function or stability in selected proteins (***Nielsen et al., 2017; Scheller et al., 2019; Abildgaard et al., 2019; Cagiada et al., 2021***). This also holds for the 154,808 variants in the 29 proteins studied here (Fig. 1a), so that variants with large ΔΔE or ΔΔG scores tend to show low fitness (low *s*_exp_). In particular, variants for which both ΔΔG and ΔΔE values are high almost always show loss of function, while those with low scores for both usually show wild-type-like fitness. As expected and discussed above, variants that do not substantially perturb stability (low ΔΔG) can both have low and high values of *s*_exp_, because substitutions may affect function through other mechanisms than loss of stability and abundance. From the stability-conservation landscape, however, it appears that such effects are captured by the ΔΔE scores so that, even for variants with low ΔΔG, conservative substitutions (low ΔΔE) tend to be associated with with high fitness, and high ΔΔE with low fitness.

To quantify the power of ΔΔE and ΔΔG for classifying variants, we extracted the top and bottom third of variant scores (*s*_exp_) as the subsets that are most clearly associated with wild-type-like and loss-of-fitness phenotypes, respectively. Next, we divided our MAVE stability-conservation landscape into a total of 16 sectors. The normalised ΔΔE scores have evenly distributed thresholds of 0.25, 0.50 and 0.75, while the ΔΔG thresholds are at 2.0, 3.0 and 4.5 kcal/mol (Fig. 1). Inspection of the sectors confirms that extreme values of ΔΔE and ΔΔG can classify variants well into the high or low fitness categories, while moderate values have lower classification power (Fig. 1b). For example when ΔΔE < 0.25 and ΔΔG < 2.0 kcal/mol, 81% of the variants are in the high fitness category, and at the opposite end, when ΔΔE > 0.75 and ΔΔG > 4.5 kcal/mol, 93% of the variants are in the low fitness category.

More generally, the two-dimensional fitness landscapes illustrate the partial interdependency of ΔΔE and ΔΔG. Most notably, it is clear that evolution selects strongly against destabilising variants so that there are almost no cases with high values of ΔΔG and low values of ΔΔE (only 4% of the variants have ΔΔE < 0.25 and ΔΔG > 3.0 kcal/mol). Also, as discussed above, low values of ΔΔG maybe associated both with high and low values of *s*_exp_, whereas variants with larger values of ΔΔG tend to have low *s*_exp_. Thus, focusing on the stable variants (ΔΔG < 2.0 kcal/mol) and only the top/bottom third of the fitness scores, we find that 38% of these variants have ΔΔE > 0.5 (Fig. 1a) and are thus more likely to be non-functional. Focusing on the variants that the GEMME analysis suggest are incompatible with what has been observed through evolution (ΔΔE > 0.5), and thus are more likely to be non-functional, we find that 47% are predicted to be unstable (ΔΔG > 2.0 kcal/mol; Fig. 1b). Thus, in line with our previous analysis (***Cagiada et al., 2021***), these results suggest that approximately half of the non-conservative substitutions are selected against due to stability effects.

### Training and Benchmarking Random Forest Models

Having established that there is an overall relationship between calculated values of ΔΔG and ΔΔE and experimental variant effects (*s*_exp_) and that both independently contribute valuable information (Fig. 1) we decided to train a machine learning model to predict variant effects from ΔΔG and ΔΔE. Before doing so, we analysed how well the data generated by the individual MAVEs correlate with the calculated values of ΔΔG and ΔΔE (Fig. 2a).

**Figure 2.**
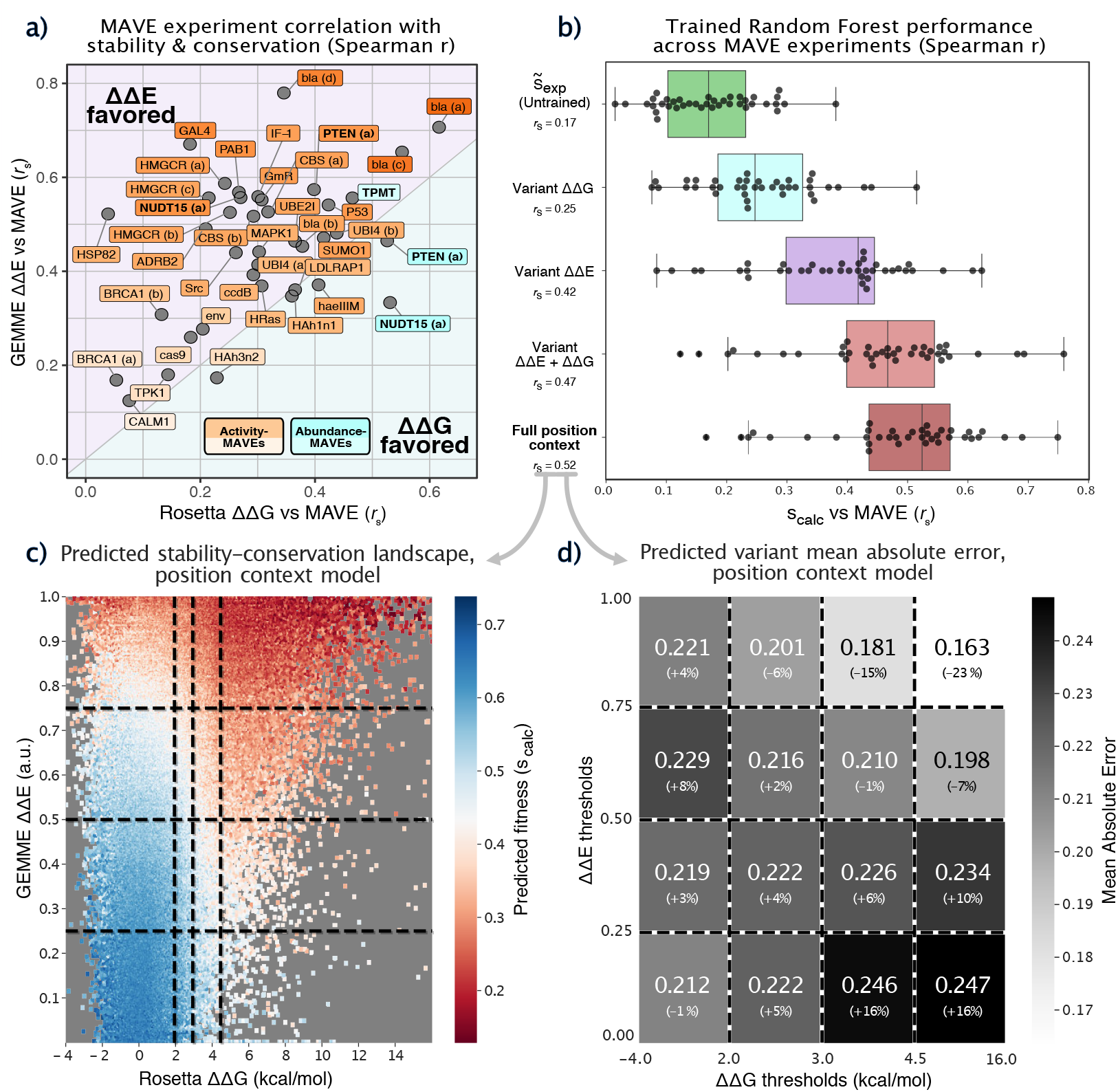
Correlations and predictions of variant effects. a) Spearman correlation coefficients between GEMME ΔΔE or Rosetta ΔΔG, and the experimental variants effects (*s*_exp_) for each of the 39 MAVEs. b) Correlation between predicted and experimental variant effects from a baseline 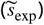 and four different machine learning models. Apart from the first row, each of the rows correspond to one of four different random forest models that used different features as input (indicated by the labels and described in detail in the main text and Methods). For each class of model, each point corresponds to one of the 39 sets of data, and the correlation coefficients were calculated in a *leave-one-protein-out* cross-validation across 29 proteins and 39 MAVE data sets. c) Predicted fitness landscape in leave-one-protein-out cross-validation of the full position context random forest model. d) Mean absolute error (MAE) and relative difference from the mean MAE from the position context random forest model.

We find a broad range of Spearman correlation coefficients (*r_s_*) between the MAVE scores (*s*_exp_) and our calculated ΔΔG and ΔΔE scores, in line with previous observations of considerable variation in correlation with variant effect predictors (***Livesey and Marsh, 2020***). Correlations range from low (*r_s_* ~ 0.1 on BRCA1 and Calmodulin-1 (CALM1) for both ΔΔE and ΔΔG) to relatively high (*r_s_* ~ 0.6 - 0.8) when predicting multiple independent MAVEs on *β*-lactamase (bla) (Fig. 2a). Overall and as expected, experimental variant effects that correlate well with ΔΔG also correlate well with ΔΔE. In line with the results shown above (Fig. 1) and previous analyses (***Nielsen et al., 2017; Abildgaard et al., 2019; Jepsen et al., 2020; Livesey and Marsh, 2020; Gerasimavicius et al., 2020; Cagiada et al., 2021***), we find that variant effects tend to be more strongly correlated with ΔΔE than ΔΔG. Two notable outliers to this observation are the abundance-based MAVEs (VAMP-seq) for PTEN (***Matreyek et al., 2018***) and NUDT15 (***Suiter et al., 2020***), which are more strongly correlated with our analysis of protein stability than conservation (labelled as PTEN (a) and NUDT15 (a) in Fig. 2a) (***Cagiada et al., 2021***). Interestingly, the VAMP-seq data for the protein TPMT (***Matreyek et al., 2018***) is slightly more strongly correlated with ΔΔE than ΔΔG. PTEN and NUDT15 variants have also been assayed for their respective biochemical functions (***Mighell et al.,2018; Suiter et al., 2020***), and the resulting *s*_exp_ scores correlate better with ΔΔE and less well with ΔΔG than the corresponding protein abundance scores (the MAVEs of respective biochemical function are labelled as PTEN (b) and NUDT15 (b) in Fig. 2a) (***Cagiada et al., 2021***). While the reasons for the low correlation between ΔΔE and ΔΔG with several of the experiments remain unclear, possible explanations include inaccuracies in stability calculations, poor sequence alignments, or experimental assays that probe properties not directly related to stability or an evolutionary-conserved function of the protein.

Next we constructed a simple baseline model that captures effects of substituting each of the 20 protein-coding amino acids for another by averaging over the normalised scores from all variants in our data set (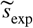; Fig. S1). Similar to previous observations (***Gray et al., 2017***) this substitution matrix captures well-known biochemical patterns. In particular we observe that substitutions of hydrophobic residues—often found in the protein core—with charged or polar residues on average lead to a large loss of fitness, while changes from a polar to a hydrophobic residue on average does not cause a substantial loss of fitness. Nevertheless, although 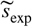 captures such chemical intuition it lacks information about structural and sequence context, and thus is overall a poor predictor of experimental variant effects (mean *r_s_* = 0.17; Fig. 2b, green).

With the experimental and computational variant data we proceeded to train a set of machine learning models to predict *s*_exp_ from our set of available ΔΔG, ΔΔE and 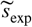 scores. We chose to use random forest models because of their robustness to outliers and noise, minimal need to adjust model hyperparameters, as well as the possibility to easily extract information about the extent to which the different features are used in the model decision process (***Breiman, 2001; Bernard et al., 2009***). We term the values predicted by the model *s*_calc_.

We used a leave-one-protein-out procedure for training, selecting one protein for validation, and training on the normalised MAVE, ΔΔE and/or ΔΔG data from all other proteins in our overall set. Thus, when more than one set of experiments had been performed on a single protein, we excluded the extra data sets during training. We assess the model by calculating the correlation between *s*_calc_ predicted from the resulting random forest with the *s*_exp_ values from all MAVEs on the protein that was left out in training, looping over all proteins one at a time.

First, we trained three random forest models using as inputs only the variant ΔΔG, only the variant ΔΔE, or both. In line with the variation in the correlation to the input data, we observe a range of correlation coefficients from these models, with the combined ΔΔG and ΔΔE model correlating with the normalised MAVE scores with a median *r_s_* = 0.47 (Fig. 2b, orange). The ΔΔG-only model (*r_s_* = 0.25, Fig. 2b, cyan) only performs slightly better than 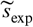 (green), and the ΔΔE-only model (*r_s_* = 0.42, Fig. 2b, purple) is again closer to capturing the experimental outcomes. We note here that several of the correlation coefficients observed in these analyses (Fig. 2b) are smaller than those obtained when correlating directly with the *s*_exp_ values (Fig. 2a). This is because we in the random forest models aim to capture the relationship between e.g. ΔΔG and *s*_exp_ using a single model and scale, whereas the correlations in Fig. 2a effectively correspond to 39 distinct ‘models’.

As variant effects depend both on the specific substitutions but also the context within the protein, we trained a more complex’position-context’model that also takes into account the scores for other substitutions at the given protein position. Specifically, in addition to the ΔΔE and ΔΔG value for the variant to be predicted, we input the entire set of 20 ΔΔG and 20 ΔΔE values at a position, representing the stability and conservation scores for all single amino-acid variants at the given position, as well as the mean score for these. We further add three features from the baseline model, 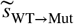 (the average score when mutating from the wild-type to the specific variant (mutant) residue), 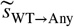 (the average score for substitutions from the specific wild-type amino acid to any of the other 19), and 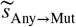 (the average score when mutating each of the 19 other amino acids to the specific variant amino-acid). The resulting position-context random forest model yields an improved performance (median *r_s_* = 0.52, Fig. 2b, brown). The model also successfully recapitulates the trends in the stability-conservation landscape (Fig. 2c and Fig. S2). Looking more closely at how the model performs in different sectors of this landscape, we find that predictions generally have a similar mean absolute error (MAE) in most sectors (Fig. 2d and Fig. S3) with MAE ≈ 0.22, with somewhat more accurate predictions for highly destabilizing (ΔΔG > 3.0) and non-conservative (ΔΔE > 0.50) substitutions (MAE 0.16-0.21).

### Role of Stability and Conservation in the Predictions

With the final model in hand and having shown that it recapitulates the experimentally observed stability-conservation landscape relatively well (Fig. 1a, Fig. 2c and Fig. S2), we proceed to analyse the properties of the model. Looking at the predictions of the >150,000 variant effects from the leave-one-protein-out model, we find that the position-context model predicts fitness outcomes with a Spearman correlation coefficient of 0.50 (Fig. 3a). The most dense region has a sigmoidal shape, with predictions of the experimentally-derived *s*_exp_ range 0.5-1.0 seeming largely indistinguishable by the model. The non-linear relationship is presumably an effect of the use of the mean square error between *s*_exp_ and *s*_calc_ as target when training the random forest model.

**Figure 3.**
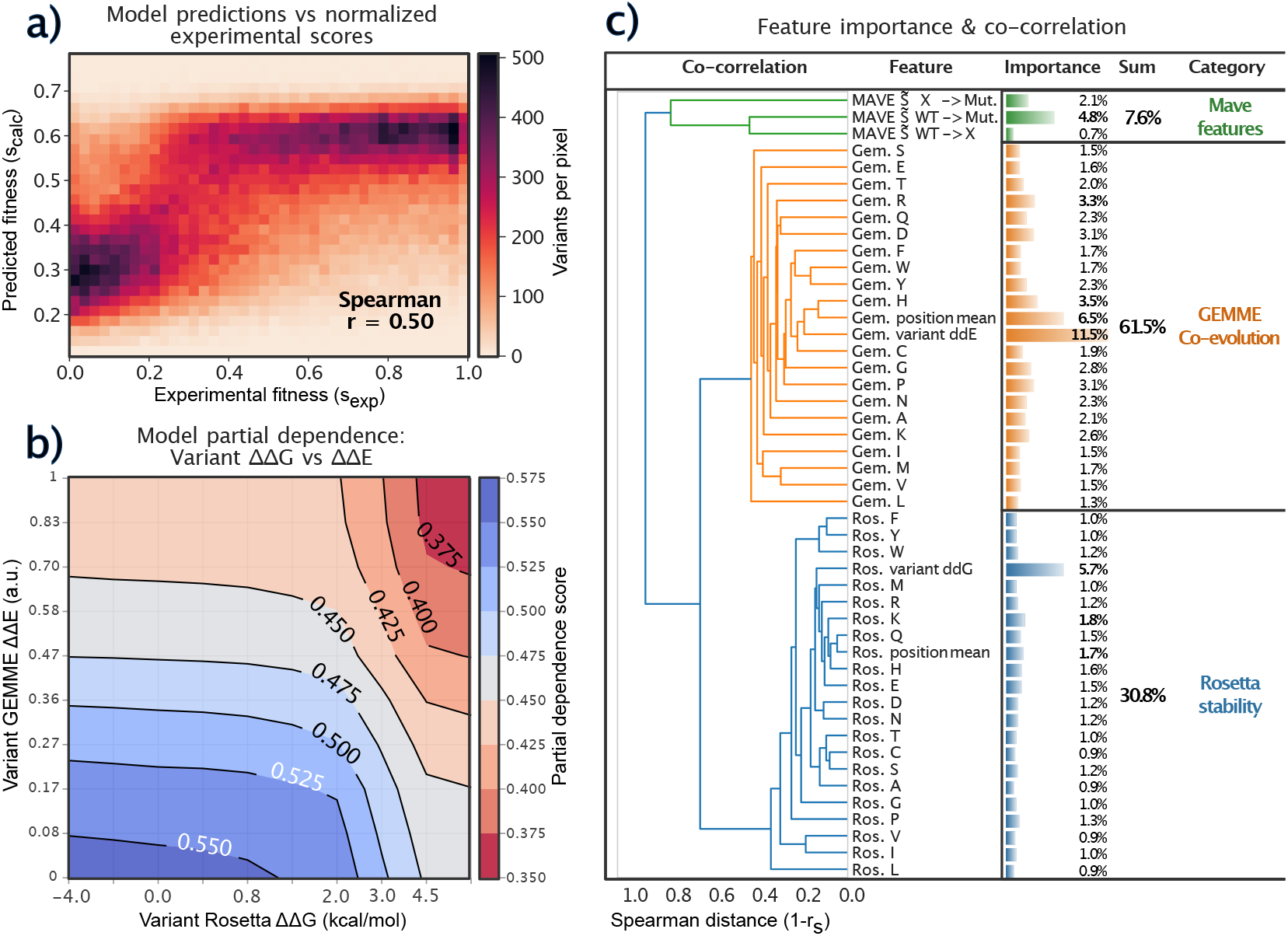
Interpretation of the position-context random forest model a) Density plot of predicted variant scores vs. normalized experimental fitness scores in leave-one-protein-out cross-validation. b) Partial dependence plot for the position-context model trained on all variants. Most of the variation in fitness scores is explained along the GEMME ΔΔE axis, with Rosetta ΔΔG only impacting variant scores at ΔΔG ≳ 2.0. Colours indicate the predicted partial score for the given simulated variant ΔΔG and ΔΔE score sampled across the entire data set, but does not take into account effects from the remaining features. Note the non-linearity of the scale of the axes. c) Position context model feature importance and dendrogram.

To gain further insight into the relative importance of the stability and conservation parameters for predicting the variant scores, we calculated the partial dependencies in the position-context random forest model. These partial dependencies quantify how the model’s predictions of *s*_calc_ depend on the chosen features, here the ΔΔG and ΔΔE for the specific variant, marginalizing over the remaining features in the model (***Molnar, 2019***). For variants that at most cause a modest change in stability (ΔΔG < 2.0 kcal/mol), the predictions are most strongly influenced by ΔΔE. This observation is in line with the notion that for stable variants, ΔΔE is a relatively good predictor of *s*_exp_ (Fig. 1a). In contrast, for more destabilizing variants (ΔΔG > 3.0 kcal/mol) the partial dependence vary with both ΔΔG and ΔΔE, in line with the finding that both of these are useful quantities to help predict variant effects in this part of the stability-conservation landscape (Figs. 1 and 2).

Wethen proceeded to examine the impact of all features in the random forest model. We stress that this so-called ‘feature importance’ only quantifies how much each feature is used overall in the prediction of *s*_calc_, but does not directly describe how and when these features are used (***Strobl et al., 2007***). We calculated the correlation between each pair of features and used these to cluster and build a dendogram of the features (Fig. 3c and Fig. S4). The features fall into three overall categories corresponding to ΔΔG, ΔΔE and 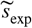.

We here remind the reader that the model uses stability and conservation effects of all possible substitutions at a specific position even when predicting the effect of a specific variant. Thus, for example when predicting the effect of a specific isoleucine to alanine substitution, the model will use as features the effect of changing that isoleucine residue to all other 19 protein-coding amino acids. As expected, however, the feature-importance calculations show that the model has its largest contribution from the specific substitution, with other substitutions playing a smaller role (Fig. 3c). In addition to using information about the specific substitution, the average ΔΔE score also has a large feature importance, which we take to mean that the model uses this to determine whether a specific position is overall restrictive in the types of substitutions that have been seen during evolution. In addition to these effects, substitutions to positively charged amino acids (e.g. ΔΔE to arginine and histidine and ΔΔG to lysine) have greater than average contributions, and we speculate that these values provide additional structural context on whether e.g. a position is buried or not (as substitutions to large positively charged residues are generally disfavoured at buried positions). We note that effects of substitutions to histidine and asparagine have previously been shown to correlate most strongly with other substitutions at the same position (***Gray et al., 2017***).

Adding up the individual contributions of the features from the three classes (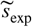, ΔΔE and ΔΔG) we find that the ΔΔE features have roughly twice the ‘importance’ as ΔΔG features, with the total importance being substantially greater than from 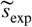 (Fig. 3c). Looking at the values for the specific variant predicted, the values are 11.8%, 5.7% and 4.8% for ΔΔE, ΔΔG and 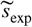, respectively, though we note caveats in interpreting these numbers very precisely (***Strobl et al., 2007***). Thus, in line with analysing simpler models using only a subset of the features (Fig. 2b) this analysis shows that all three classes of features contribute to the performance of the final model, with ΔΔE carrying the greatest weight.

### A Global Map of Variant Effects

The results above further support previous observations that analyses of sequence conservation are overall more informative than stability calculations to predict variant affects as probed by most MAVE experiments (***Jepsen et al., 2020; Livesey and Marsh, 2020; Gerasimavicius et al., 2020; Cagiada et al., 2021***). Nevertheless, they also show that stability calculations can help improve overall prediction accuracy and provide mechanistic insight into the relative roles of stability and conservation in explaining variant effects. Specifically, as argued previously (***Stein et al., 2019; Jepsen et al., 2020; Cagiada et al., 2021***), we can use ΔΔE calculations as a proxy for capturing a broad range of effects on biological function, and ΔΔG as the subset that involves specifically stability and abundance.

We thus used the data that we collected to create global maps of variant effects, and analysed these in terms of fitness, stability and conservation. Inspired by previous work (***Dunham and Beltrao, 2020***), we visualized the 6500 amino acid positions in our data set with more than 15 values of *s*_exp_ and corresponding values of ΔΔG and ΔΔE (Fig. 4). Specifically, we used UMAP dimensionality reduction (***McInnes et al., 2018***) to represent all positions in a two-dimensional map where positions with similar profiles of *s*_exp_, ΔΔG and ΔΔE are located close to one another (see Methods). Finally, we colour-coded this map using the predicted and calculated position-averaged fitness scores, the position-averaged values of ΔΔG and ΔΔE, values for 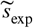 and the relative solvent accessibility of the position (RSA) (Fig. 4).

**Figure 4.**
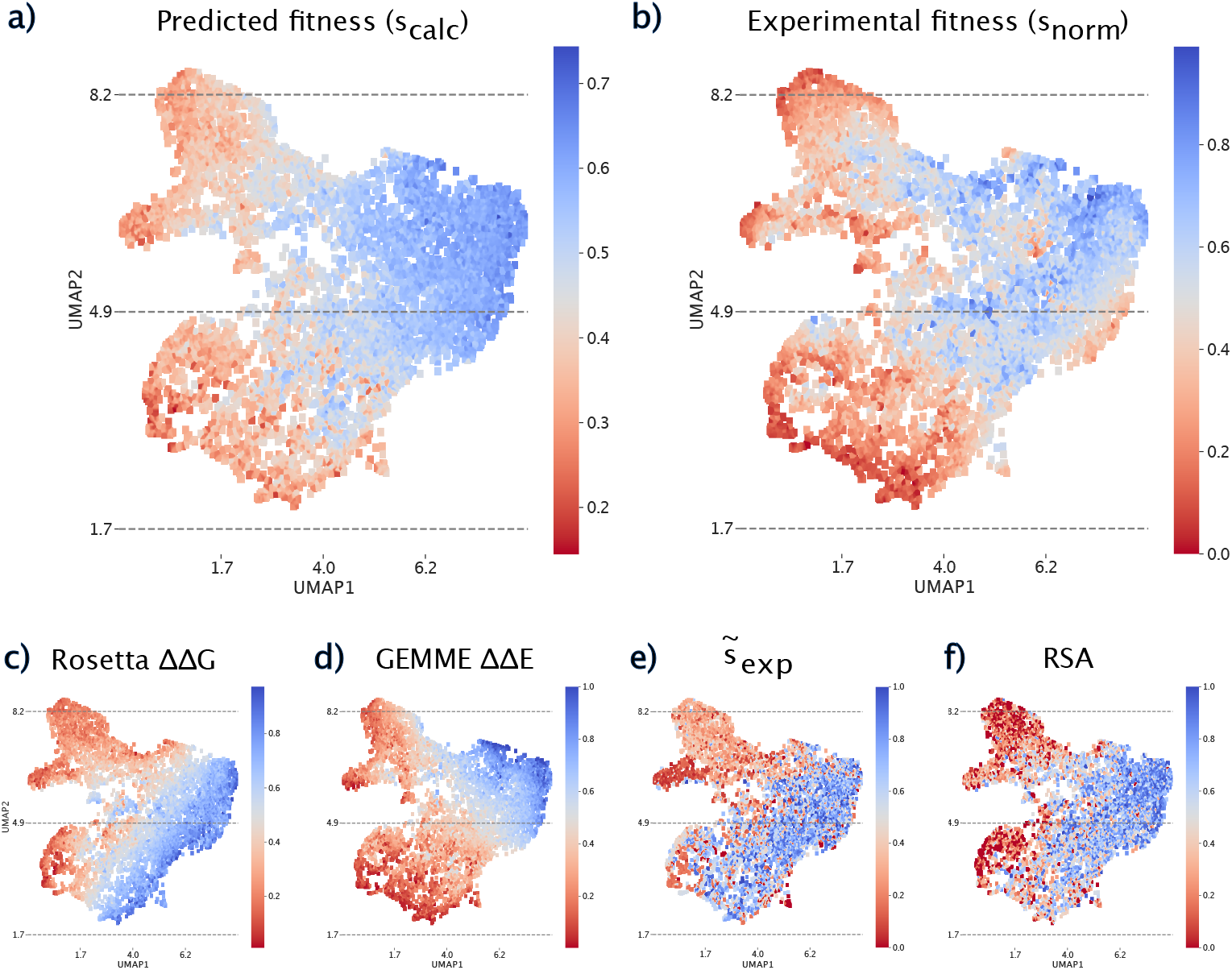
UMAP projection of over 6500 positions with data for at least 15 variants in the MAVEs. The maps are coloured by different properties and are locally-averaged using a convolutional kernel (see Methods). All scores were normalized between 0 and 1 to ease comparisons. a) Position-averaged predicted variant scores from the position-context random forest model. b) Position-averaged experimental fitness scores. c) Position-averaged ΔΔG values. d) Position-averaged ΔΔE values. e) 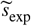. f) Relative solvent accessibility from protein structure.

As expected from the fact that the model was trained to predict the experimental data, the map coloured by *s*_calc_ (Fig. 4a) and *s*_exp_ (Fig. 4b) closely resemble one another, though with a few differences highlighting the imperfection of the model. In particular, the maps reveal two regions (top left and bottom left of the maps) that are enriched in low-fitness variants. The map of stability effects (Fig. 4c) show a gradual change moving from the top left to the bottom right, and illustrates that in particular the top left part of the map corresponds to positions where variants on average are destabilizing. In contrast, the map coloured by evolutionary scores (Fig. 4d) reveal a gradient from the bottom left to the top right, and identifies many of the same regions as the experimental map in terms of low scores. Comparing the maps coloured by the ΔΔG and ΔΔE scores reveal two regions (top left, and part of the bottom left) with amino acid positions where it appears that variants cause loss of function due to loss of stability. Comparison with the map indicating solvent accessibility (Fig. 4f) shows clearly that many of these positions are buried inside the protein structure. The same comparisons also show that a group of positions near the bottom are enriched in positions where variants lose function in ways that are not due to stability, but that can nonetheless be discovered using analyses of the evolutionary record. Finally, looking at the map coloured by the average 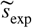 scores (Fig. 4e) shows similarities to both the stability map (Fig. 4c) and solvent accessibility map (Fig. 4f), in line with the fact that these are buried positions and that 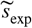 captures basic physico-chemical aspects of amino acid chemistry and protein structure (Fig. S1).

## Discussion

We have analysed the relationship between experimental measurements of variant effects on protein function and computational analysis of protein stability and conservation. We collected over 150,000 measurements from multiplexed assays of variant effect in 29 proteins, and compared them with predictions of changes in protein stability (ΔΔG) and evolutionary conservation (ΔΔE). Our goal was two-fold. First, we aimed to examine how well these computed scores could predict variant effects, and second we wanted to shed further light on how often changes in protein stability may perturb protein function.

In general, and in line with previous observations, we find that our analysis of conservation of sequences through evolution (ΔΔE) is more strongly correlated with the experimental measurements than predictions of changes in thermodynamic stability (ΔΔG) (Fig. 2a). We note here that although each of the 39 experiments provides a systematic and comprehensive analysis of variant effects, they were conducted using very different assays for selection/screening and in different organisms. Together with uncertainties in the experimental and predicted values, this likely explains the substantial variation that we observe in agreement between experimental scores and computed values. Thus, we defer in general from analysing individual proteins and variants and focus on the global agreement. As also previously observed (***Cagiada et al., 2021***), the VAMP-seq experiments that specifically probe protein abundance tend to be more strongly correlated with stability effects than measurements probing protein activity. Finally, we note that while discrepancies between experimental measurements of variant effects and computational predictions can point to shortcomings in the prediction models, these may also reveal aspects of the protein’s function that the assay was not sensitive to.

Our global stability-conservation landscape (Fig. 1a) reveals the interdependency of stability and conservation. As evolution tends to disfavour unstable proteins, we find that variants with high ΔΔG tend to have large values of ΔΔE and generally low fitness (low *s*_exp_). The reverse relationship, however, is not true because being stable is a necessary, but not sufficient, criterion for being functional. Examining the stable variants (low ΔΔG) we find that our analysis of the evolutionary record (ΔΔE) can be used to predict with reasonable accuracy whether a variant retains function or not (Fig. 1b). In our recent analysis of data probing activity and abundance in NUDT15 and PTEN, we showed that about half of the variants that lose function, do so because they become unstable and are found at low abundance. In line with this, we find that roughly half of the variants that have high ΔΔE scores also have high values of ΔΔG, and hypothesize that the remaining variants (low ΔΔG, high ΔΔE) lose function because of substitutions in functionally important residues (***Cagiada et al., 2021***).

Based on the partial correlation between both ΔΔE and ΔΔG with *s*_exp_ we built a prediction model with these parameters as input. The final position-context model reaches an accuracy that surpasses that of models that solely use ΔΔE or ΔΔG (Fig. 2b). This suggests that although ΔΔE calculations to some extent capture stability effects, they can still be improved by explicitly including calculated values of ΔΔG. Indeed, the model is most accurate for variants that both ΔΔE and ΔΔG suggest would be non-functional (Fig. 2d).

The random forest model recapitulates key aspects of the global stability-conservation landscape (Fig. S2) including the interdependency of ΔΔG and ΔΔE. A more detailed analysis of the random forest model shows that ΔΔE indeed is the more informative quantity when ΔΔG is small, whereas both ΔΔG and ΔΔE contribute to accuracy when ΔΔG is large (Fig. 3b). This is also reflected in the feature importance analysis which shows that overall the various ΔΔE terms contribute roughly twice the importance of the ΔΔG terms among the decision trees (Fig. 3c). Similar effects are also seen in our global maps of variant effects (Fig. 4), which reveal an almost orthogonal gradient of scores for ΔΔG and ΔΔE, and indicate how the combination of these two maps contribute to the final predicted scores. We also show that including contextual information about effects of other substitutions at the same position further increases correlation with experimental scores. We speculate that this context captures additional information about the structural, biochemical and functional requirements at this position.

There are several possibilities for future improvements of the prediction methods used here. First, a number of methods have recently been developed to analyse sequence information (***Riesselman et al., 2018; Alley et al., 2019; Frazer et al., 2020; Hsu et al., 2021***) and could be tested instead of the GEMME method that we used. Second, while the Rosetta method that we used to predict ΔΔG is among the most accurate methods for stability prediction, it is computationally expensive and requires access to protein structures. Thus, future work is needed on assessing the utility of predicted structures (both from template-free and template-based methods), as well as testing and developing more computationally efficient methods for predicting stability effects.

Another area for further development is to analyse and improve predictions in areas of intermediate values of ΔΔG and ΔΔE, where errors in the predicted values would have a greater impact. While our results overall conform to the expected relationship between ΔΔE and ΔΔG, more subtle effects can modulate this relationship. For example, it has been observed that active site residues may actually be destabilizing (***Shoichet et al., 1995***), and more detailed stability calculations may thus aid in detecting such effects.

The types of analyses presented here may also be used to improve or interpret experiments. For example, when constructing an experimental assay it might be useful to compare experimental variant effects to calculations of ΔΔE to examine whether the assay is sensitive to evolutionarily-conserved functions. Also, it has recently been demonstrated that biophysical ambiguities prevent accurate predictions of how substitutions combine to affect phenotype (***Li and Lehner, 2020***), and we suggest that biophysical models and predictions of variant effects may help alleviate some of these ambiguities. Finally, a random forest model combining sequence and structural features has been used to predict pathogenicity of human missense variants (***Ponzoni et al., 2020***).

In summary our large scale analysis contributes to the mounting evidence for the important role for loss of stability in loss of function. In addition to the improvements observed when combining conservation analysis with structure-based stability predictions, our analyses also help pinpoint those variants that lose function due to stability. Such information helps provide mechanistic insight into specific variants and proteins, but may also be a starting point for developing therapies. In particular, variants that lose function due to loss of abundance, but whose intrinsic function is not otherwise perturbed, would be ideal targets for approaches that aim to restabilize or otherwise restore protein levels (***Balch et al., 2008; Kampmeyer et al., 2017; Stein et al., 2019; Henning et al., 2021***).

## Methods

### ΔΔG calculations using Rosetta

We searched the Protein Data Bank (***Rose et al., 2010***) using the BLASTp webserver (***McEntyre J, 2002***) with default settings using as query the sequences for which the MAVE was performed. If only a part of a protein is covered by mutagenesis data, we only searched for structures of that part. We chose structures by balancing highest available coverage and resolution, selecting structures solved using X-ray crystallography when available. All calculations were carried out using the Rosetta version with GitHub SHA `28f338acfb3bfd87048b38a04772486975dc83fa` from July 2, 2020. We first relaxed the structures using the relax application and the following flags:

~~~
-fa_max_dis 9
-relax:constrain_relax_to_start_coords
-ignore_unrecognized_res
-missing_density_to_jump
-nstruct 1
-relax:coord_constrain_sidechains
-relax:cartesian
-beta
-score:weights beta_nov16_cart
-ex1
-ex2
-relax:min_type lbfgs_armijo_nonmonotone
-flip_HNQ
-no_optH false
~~~

Subsequently, we carried out saturation mutagenesis to calculate ΔΔG for each single amino acid substitution using the Cartesian ΔΔG protocol and the `beta_nov16_cart` energy function with three iterations as previously described (***Park et al., 2016; Frenz et al., 2020***). Flags for the ΔΔG calculations were:

~~~
-fa_max_dis 9.0
-ddg::dump_pdbs false
-ddg:iterations 3
-score:weights beta_nov16_cart
-missing_density_to_jump
-ddg:mut_only
-ddg:bbnbrs 1
-beta_cart
-ex1
-ex2
-ddg::legacy true
-optimize_proline true
~~~

Scores from the three iterations were averaged. Values of ΔΔG in Rosetta Energy Units were divided by 2.9 to bring them onto a scale corresponding to kcal/mol (Frank DiMaio, University of Washington; personal correspondence;***Jepsen et al. (2020)***).

### Evolutionary conservation analysis using GEMME

We calculated evolutionary conservation scores (ΔΔE) using GEMME, a global epistatic model for predicting mutational effects (***Laine et al., 2019***), based on a multiple sequence alignment of homologs of each protein of interest. We used the sequence used in the MAVE experiment as input to HHblits (version 2.0.15 and settings `-e 1e-10 -i 1 -p 40 -b 1 -B 20000`)to search `UniRef30_2020_03_hhsu` (***Mirdita et al., 2017; UniProt Consortium, 2019; Steinegger et al., 2019***). We de-gapped the alignment with respect to the MAVE sequence, removed sequences with 50% gaps and used the output alignment as input to GEMME, with default settings. Finally, we rank-normalized the output ΔΔE scores and scaled them to a [0,1] scale.

### Collecting and normalizing data from MAVEs

We downloaded 39 data sets that had been generated by MAVEs from publicly available repositories including MAVEdb (***Esposito et al., 2019***) and a compilation by ***Livesey and Marsh (2020)*** (see Tables S1 and S2). We used the variant fitness scores as presented in the original publication including possible normalization. Three data sets (P53, MAPK1 and Src) showed a reverse relationship between variant scores and ΔΔG and ΔΔE and were therefore reversed to match our convention of high scores for wild-type-like activity and low scores for low activity. We then rank-normalised and scaled these scores to a [0,1] range. We term these normalized scores *s*_exp_.

We next merged each data set from the MAVEs with the corresponding ΔΔE and ΔΔG scores aligning the sequences based on using Biopython, `pairwise2.align.globalds(target_seq.upper(), seq.upper(), MatrixInfo.blosum62, −3, −1)`.

### Baseline substitution model

For each MAVE data set, we calculated a 20 × 20 amino-acid matrix containing the average of each the 400 possible Wild-type – Mutant variant scores (e.g. the average *s*_exp_ for all valine to alanine substitutions). We then calculated a global substitution score matrix 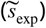 by averaging these matrices.

### Features and Random Forest models

For each variant, we extracted in total 47 computational features. Depending on the model, each *s*_exp_ was matched with its variant ΔΔG and ΔΔE score, as well as the 20 ΔΔG and 20 ΔΔE values corresponding to all available scores at the respective position, plus the mean ΔΔG and ΔΔE scores for the position. We note that this results in some redundancy in the use of the data, but simplifies the data structure in the model. We also included three global features from the baseline substitution model: (i) The mean MAVE score 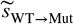, (ii) 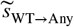 was calculated as the mean for any substitution from the given wild-type amino acid, and (iii) 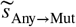 as the mean for mutating to the specified amino-acid from any wild-type. Thus, these three features correspond to (i) an entry in 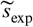 as well as the (ii) row-average, and (iii) column-average of 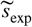 (Fig. S1).

We trained the Random Forest models using the RandomForestRegressor in Scikit-Learn (***Pedregosa et al., 2011***), with a mean-squared-error loss function, 150 trees and a minimum of 15 samples per leaf.

We first trained two models, using the variant ΔΔG or ΔΔE score only (1 feature). Next we trained a combined model using both features. Training of the position-context model was performed with all 47 available features. We iteratively trained the model and evaluated validation set performance in a leave-one-protein-out cross-validation, with the removal of all validation data sets for the selected protein from the training data set for each training round.

### Model analysis

To extract feature importance from a globally-trained model, we trained a new random forest model using all 47 features and all available data sets. The feature co-correlation dendrogram was constructed by first calculating the Spearman correlation between each pair of features, which in turn was converted into a distance as 1 — *r_s_*, and used as input to build the dendogram using SciPy (***Virtanen et al., 2020***). We measured partial dependence using PDPbox (***Jiangchun, 2018***).

### Relative solvent accessibility

We used DSSP (***Kabsch and Sander, 1983***) to calculate the relative solvent accessibility (RSA) using the structures that we also used for the Rosetta calculations. The RSA was used in analysis but not in training.

### UMAP projection of MAVE positions

To map variant scores to positions, we removed positions with fewer than 15 experimental *s*_exp_ scores from the data set, and calculated position means for *s*_exp_, ΔΔE, ΔΔG, 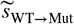 and RSA. Any missing *s*_exp_ scores were then imputed using `sklearn.impute.SimpleImputer` from the amino-acid mean across all extracted positions, and all scores were then normalized to a 0-1 range. For computational efficiency we reduced the data set to 20 features using PCA before projecting the data into two dimensions (UMAP1 and UMAP2), using `UMAP-learn` with default settings (***McInnes et al., 2018***).

### Data availability

Scripts and data to repeat our analyses are available via https://github.com/KULL-Centre/papers/tree/master/2021/ML-variants-Hoie-et-al

## Acknowledgement

Our research is supported by the PRISM (Protein Interactions and Stability in Medicine and Genomics) centre funded by the Novo Nordisk Foundation (NNF18OC0033950, to A.S. and K.L.-L.) and the Lundbeck Foundation (R272-2017-4528, to A.S.)

## Supporting Information

**Supporting Table 1.** Overview of experimental data generated by MAVE (part 1 shows 31/39 data sets). The Spearman correlation coefficients between experimental scores and the matching variant GEMME ΔΔE, Rosetta ΔΔG and substitution averages 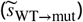 re shown.

| Name | Organism | Assay | Predicted | GEMME | Rosetta | MAVE<br>WT → Mut | PDB | MAVE reference |
| --- | --- | --- | --- | --- | --- | --- | --- | --- |
| bla (a)<br>(P62593) | E. coli | Antibiotic resistance | 0.81 | 0.71 | 0.62 | 0.48 | 1ZG4_A | Firnberg et al 2014 |
| bla (c)<br>(P62593) | E. coli | Antibiotic resistance | 0.75 | 0.65 | 0.55 | 0.49 | 1ZG4_A | Stifler et al 2015 |
| bla (d)<br>(P62593) | E. coli | Antibiotic resistance | 0.69 | 0.78 | 0.35 | 0.29 | 1ZG4_A | Jacquier et al 2013 |
| GAL4<br>(P04386) | Yeast | Two-hybrid assay | 0.66 | 0.67 | 0.18 | 0.36 | 3COQ_A | Kitzman et al 2015 |
| PAB1<br>(P04147) | Yeast | Competitive growth assay | 0.62 | 0.57 | 0.27 | 0.48 | 6R5K_D | Melamed et al 2013 |
| NUDT15 (b)<br>(Q9NV35) | Human | Drugsensitivity, FACS | 0.62 | 0.56 | 0.27 | 0.25 | 5BON_A | Suiter et al 2019 |
| TPMT<br>(P51580) | Human | Abundance, GFP fluorescence | 0.61 | 0.56 | 0.47 | 0.38 | 2H11_A | Matreyek et al 2018 |
| PTEN (b)<br>(P60484) | Human | Competitive growth assay | 0.60 | 0.57 | 0.40 | 0.36 | 1D5R_A | Mihell et al 2018 |
| P53<br>(P04637) | Human | Competitive growth assay, P53 agonists | 0.60 | 0.54 | 0.42 | 0.28 | 4QO1_B | Giacomelli et al 2018 |
| GmR<br>(Q53396 template) | E. Coli | Antibiotic resistance | 0.57 | 0.55 | 0.31 | 0.25 | 6BVC_A | Dandageet al 2018 |
| HMGCR (a)<br>(P04035) | Human | Functional complementation, atorvastatin | 0.57 | 0.59 | 0.24 | 0.32 | 1DQ8_A | Jiang et al, 2019 |
| infA<br>(P69222) | E. Coli | Competitive growth assay | 0.56 | 0.56 | 0.30 | 0.33 | 1AH9_A | Kelsic et al 2016 |
| CBS (a)<br>(P35520) | Human | Competitive growth assay | 0.55 | 0.53 | 0.32 | 0.35 | 4COO_A | Sun et 2020 |
| SUMO1<br>(P63165) | Human | Competitive growth assay, yeast | 0.55 | 0.47 | 0.42 | 0.42 | 1WYW_B | Weile et al 2017 |
| PTEN (a)<br>(P60484) | Human | Abundance, GFP fluorescence | 0.54 | 0.46 | 0.53 | 0.40 | 1D5R_A | Matreyek et al 2018 |
| HMGCR (c)<br>(P04035) | Human | Functional complementation, rosuvastatin | 0.53 | 0.56 | 0.21 | 0.29 | 1DQ8_A | Jiang et al, 2019 |
| UBE2I<br>(P63279) | Human | Competitive growth assay, yeast | 0.53 | 0.46 | 0.37 | 0.40 | 2GRN_A | Weile et al 2017 |
| HMGCR (b)<br>(P04035) | Human | Functional complementation, control | 0.53 | 0.53 | 0.25 | 0.29 | 1DQ8_A | Jiang et al, 2019 |
| bla (b)<br>(P62593) | E. coli | Antibiotic resistance | 0.52 | 0.45 | 0.38 | 0.29 | 1ZG4_A | Deng et al 2012 |
| CBS (b)<br>(P35520) | Human | Competitive growth assay | 0.52 | 0.52 | 0.29 | 0.31 | 4COO_A | Sun et 2020 |
| ADRB2<br>(P07550) | Human | Competitive growth assay | 0.51 | 0.49 | 0.21 | 0.33 | 2R4R_A | Jones et al 2019 |
| haellIM<br>(P20589) | H. Aegyptius | Competitive growth assay | 0.50 | 0.37 | 0.41 | 0.39 | 3UBT_A | Rockah-Shmuel et al 2015 |
| Src (b)<br>(P12931) | S. Cerevisae | Kinase activity, catalytic domain | 0.48 | 0.44 | 0.26 | 0.29 | 2H8H_A | Ahler et al 2019 |
| UBI4 (a)<br>(P0CG63) | Yeast | Competitive growth assay | 0.48 | 0.48 | 0.44 | 0.44 | 3OLM_D | Roscoe et al 2013 |
| MAPK1<br>(P28482) | Human | Competitive growth assay, doxycycline | 0.47 | 0.44 | 0.30 | 0.22 | 4QTA_A | Brenan et al 2016 |
| HSP82<br>(P02829) | Yeast | Competitive growth assay | 0.47 | 0.52 | 0.04 | 0.26 | 2CG9_A | Mishra et al 2016 |
| ccdB<br>(P62554) | E. coli | Reverse survival assay | 0.45 | 0.39 | 0.29 | 0.39 | 1VUB_A | Adkar et al 2012 |
| UBI4 (b)<br>(P0CG63) | Yeast | FACS-sorting | 0.44 | 0.41 | 0.30 | 0.47 | 3OLM_D | Roscoe & Bolon, 2014 |
| HA-H1N1<br>(A0A2Z5-U3Z0) | Influenza virus | Competitive replication assay | 0.44 | 0.35 | 0.36 | 0.41 | 6MYA_A | Doud & Bloom, 2016 |
| NUDT15 (a)<br>(Q9NV35) | Human | Abundance, GFP fluorescence | 0.44 | 0.33 | 0.53 | 0.37 | 5BON_A | Suiter et al 2019 |
| LDLRAP1<br>(Q5SW96) | Human | Two-hybrid assay, OBFC1 interaction | 0.43 | 0.36 | 0.37 | 0.41 | 3SO6_A | Jiang et al, 2019 |
| Hras<br>(P01112) | Human | Two-hybrid assay | 0.43 | 0.37 | 0.31 | 0.36 | 5E95_A | Bandaru et al 2017 |

**Supporting Table 2.**
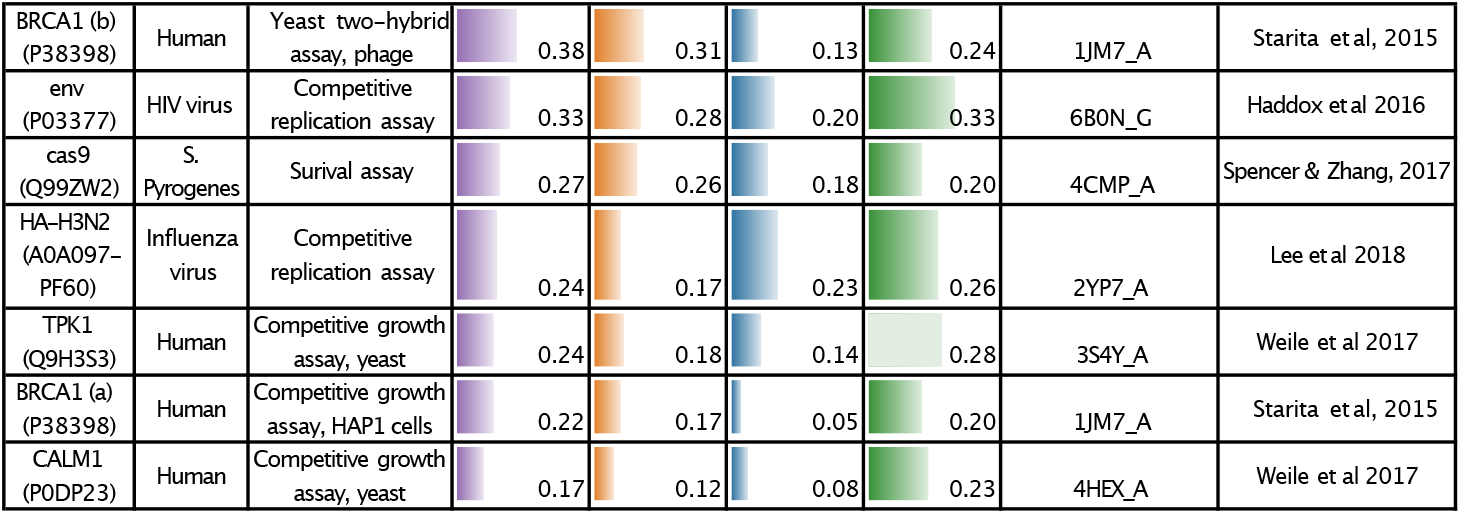
Overview of experimental data generated by MAVE (part 2 shows 8/39 data sets). The Spearman correlation coefficients between experimental scores and the matching variant GEMME ΔΔE, Rosetta ΔΔG and substitution averages 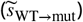 are shown.

**Supporting Figure 1.**
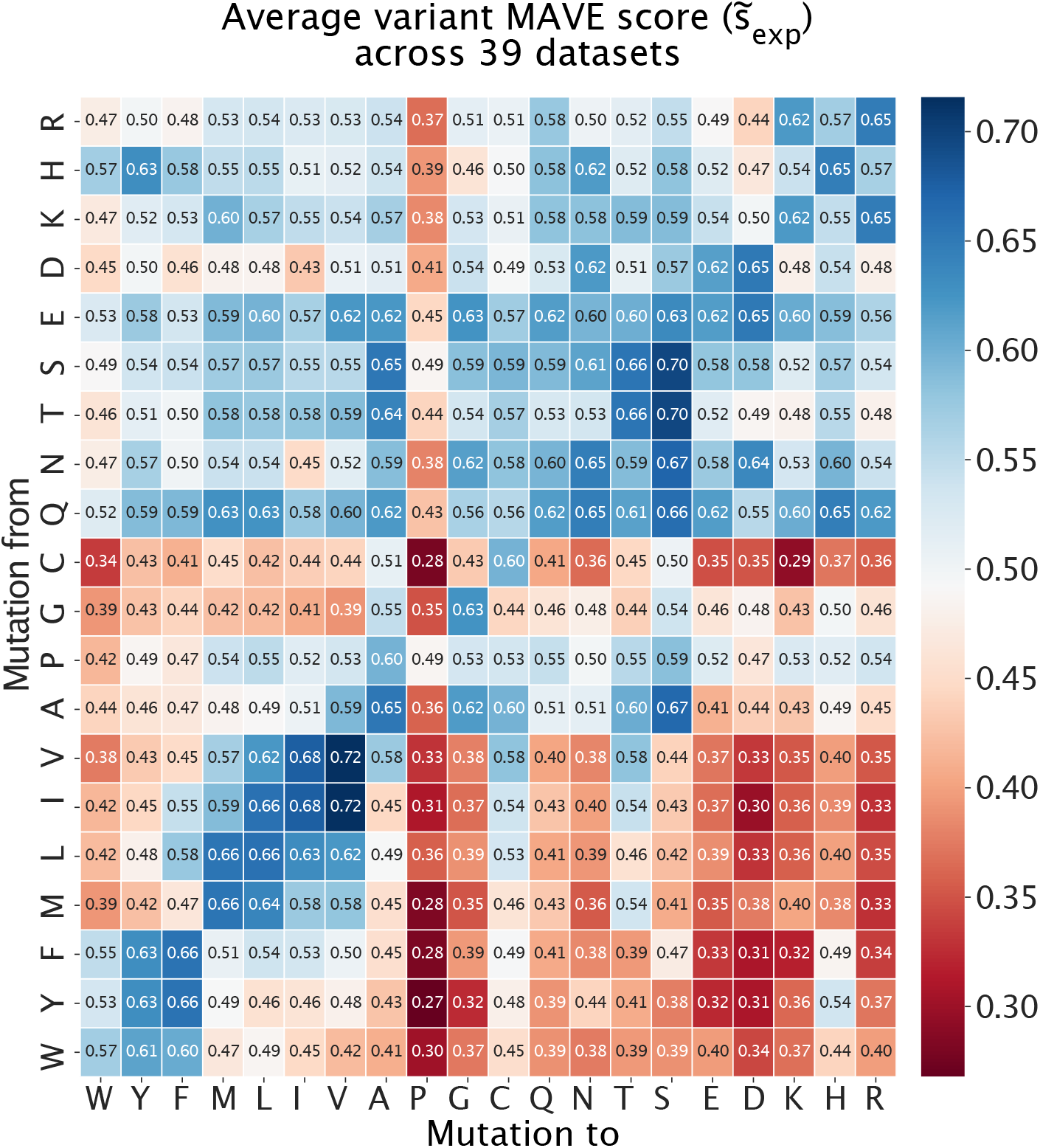
Average scores (after normalization) across 39 data sets generated by MAVEs (see Methods). Lower scores (red) indicate lower fitness, with normalized experimental scores ranging from 0-1. Substitutions from hydrophobic and typically buried residues to polar or charged residues tend to cause substantial loss of fitness. Substitutions to proline and from cysteine are (on average) also particularly disruptive. These values can be used as a baseline to predict variant effects (Fig. 2b).

**Supporting Figure 2.**
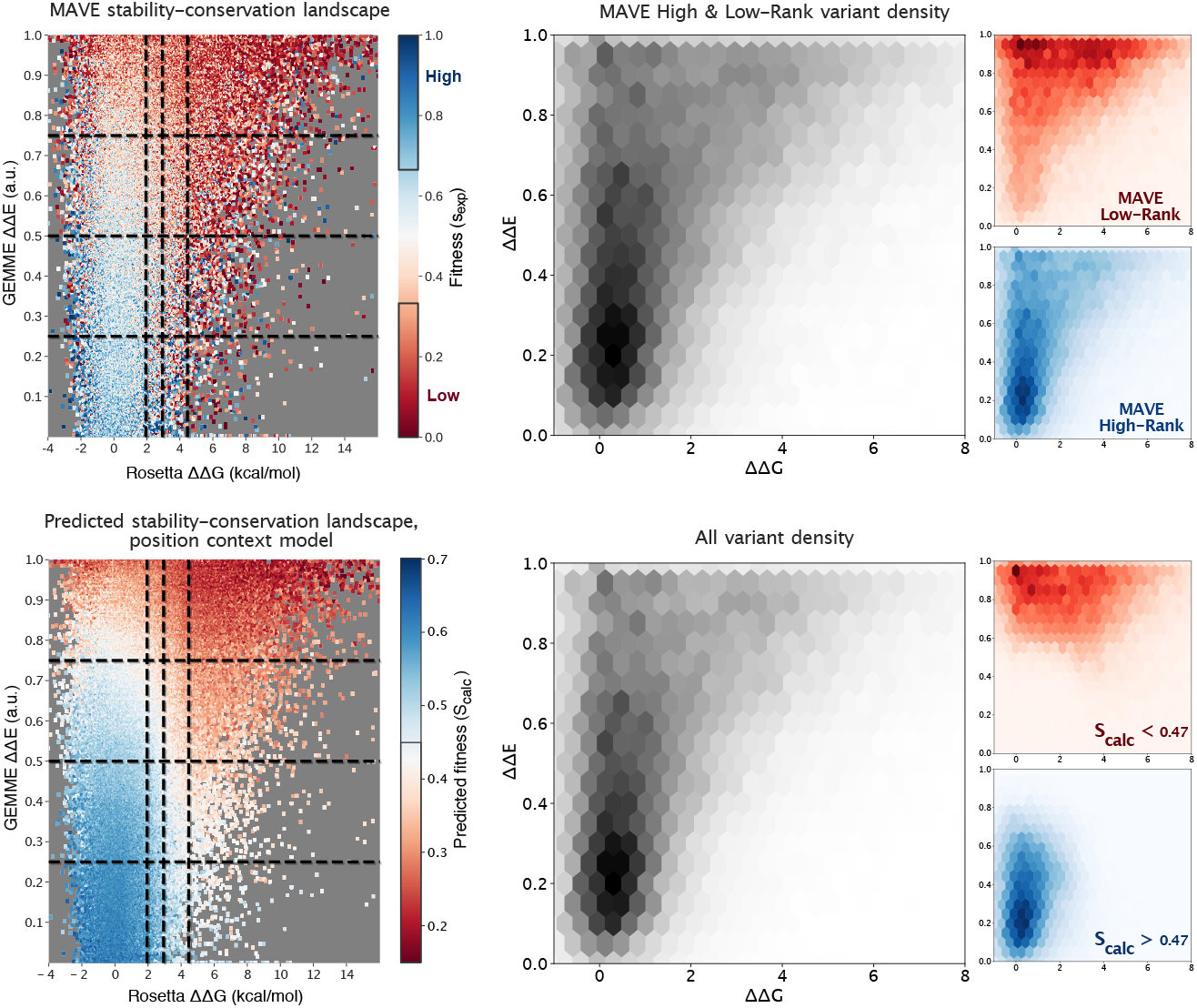
Comparison of predicted vs. normalized MAVE scores for variants in ΔΔG/ΔΔE landscape, and density of variants (High/Low only: top middle, and all variants: bottom middle). Density plots are further highlighted by isolated MAVE Low or High-Rank variants, or predicted Low or High. Predicted *s*_calc_ classification threshold is set at the median value (0.47).

**Supporting Figure 3.**
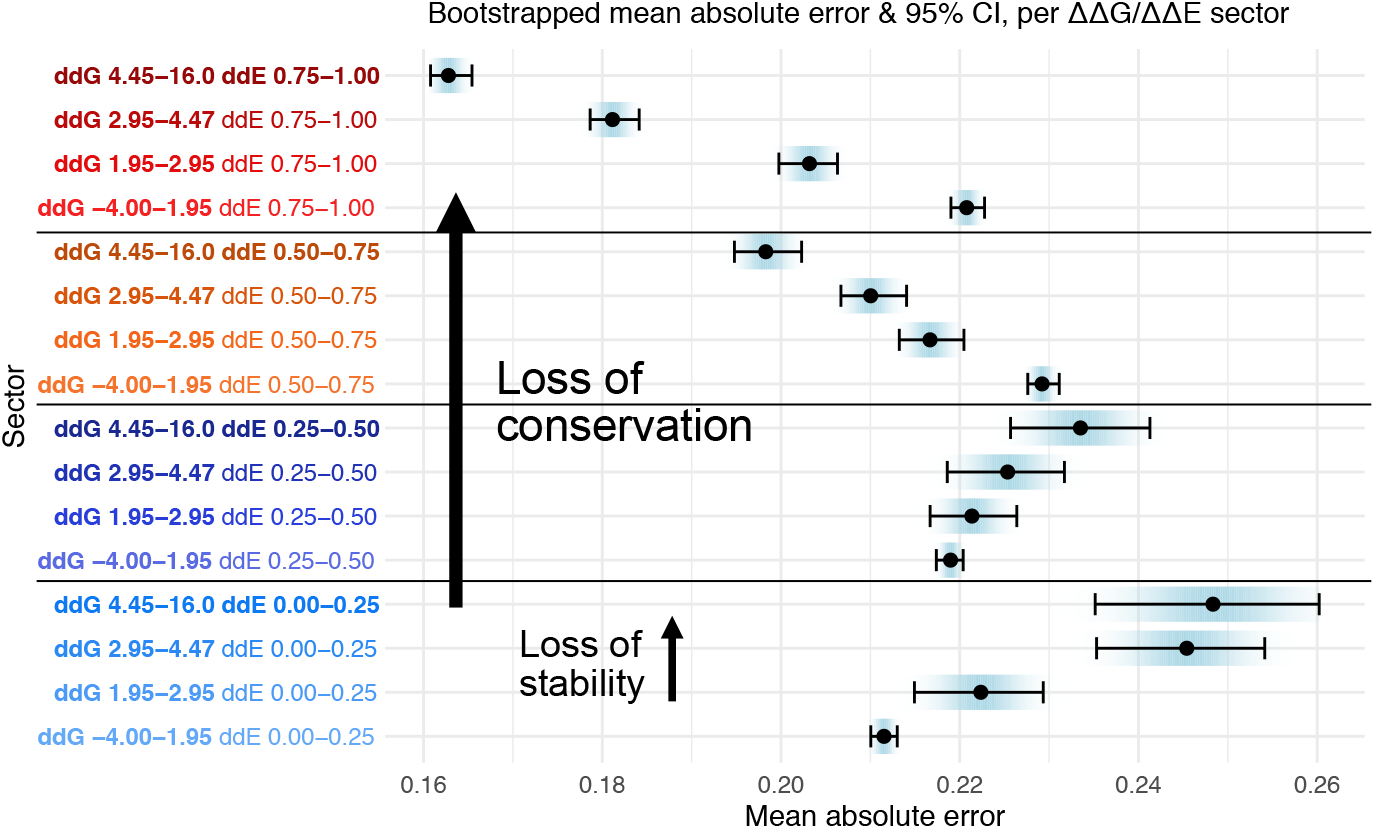
Bootstrapped mean and 95 % confidence interval for variants within each ΔΔG/ΔΔE sector of the predicted MAVE fitness landscape (Fig. 2C/D), sampled 200 times per sector. Variant loss of stability (ΔΔG) increments within each line-group, while loss of conservation (ΔΔE) increments between each line-group. High values of ΔΔG significantly improve mean absolute error for most variants with ΔΔE > 0.50.

**Supporting Figure 4.**
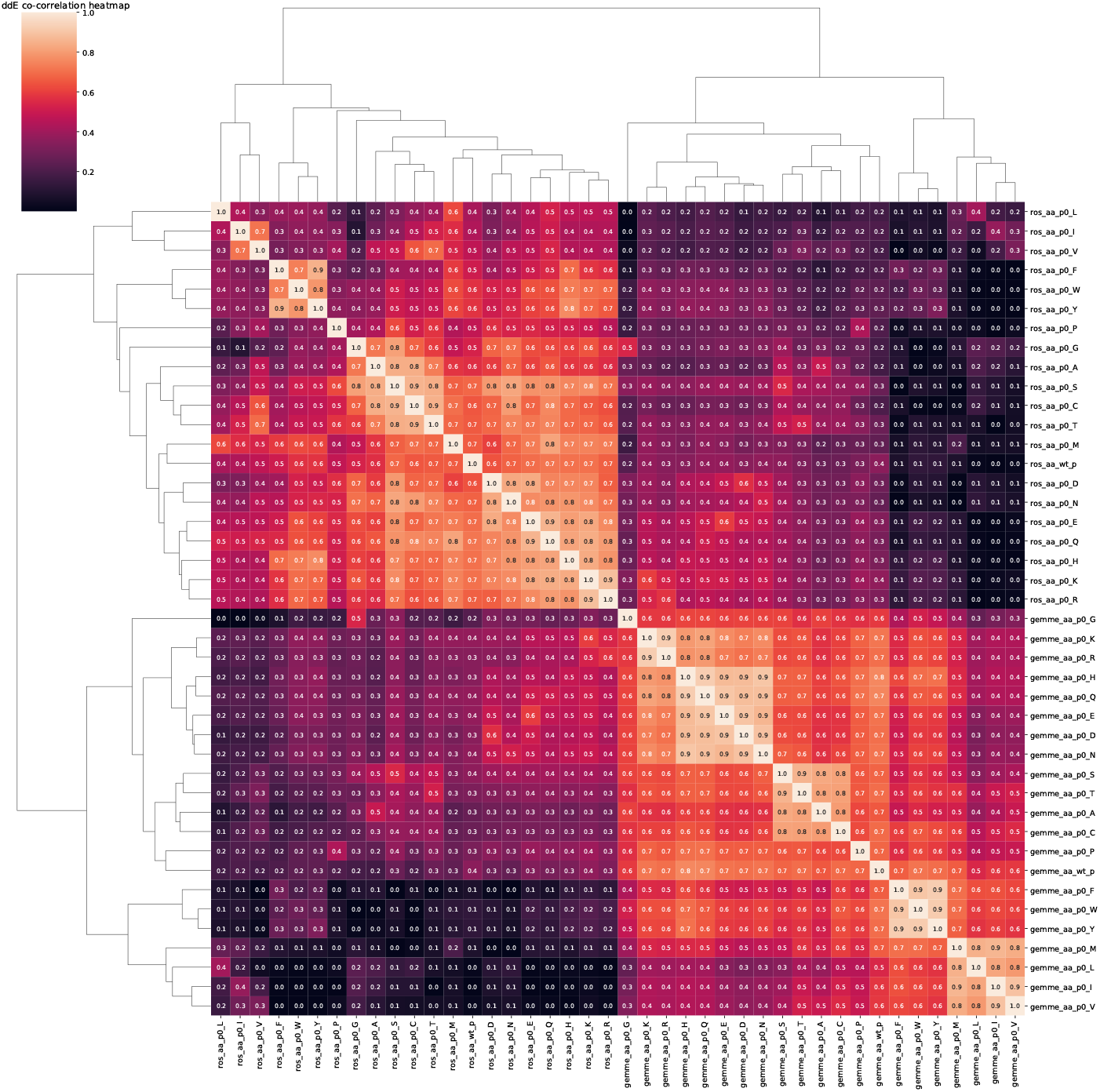
Matrix of correlations between features underlying the dendrogram in Fig. 3c

## Notes

https://github.com/KULL-Centre/papers/tree/master/2021/ML-variants-Hoie-et-al

### Competing Interest Statement

The authors have declared no competing interest.

